# The MET growth signaling complex drives Alzheimer’s Disease-associated brain pathology in aged Shugoshin 1 mouse cohesinopathy model

**DOI:** 10.1101/2024.03.26.586833

**Authors:** Chinthalapally V. Rao, Julie Crane, Ben Fowler, Yuting Zhang, Hiroshi Y. Yamada

## Abstract

The understanding on molecular processes toward Late-onset Alzheimer’s Disease (LOAD) has been insufficient to design LOAD intervention drugs. Previously, we discovered transgenic genomic instability model mice Sgo1-/+ accumulate cerebral amyloid-beta in old age. We proposed the “amyloid-beta accumulation cycle” hypothesis, in which cytotoxic, mitogenic and aneuploidgenic amyloid can create an autonomous mitotic cycle leading to accumulation of itself. However, the nature of the growth signaling that drives cells toward pathogenic mitotic cycle remained unidentified. In this study, we hypothesized that the aged Sgo1-/+ mice brains would show signs of mitogenic signaling activation, and searched for growth signaling activated in the vicinity of amyloid-beta, with spatial analysis on the cortex and hippocampus of Sgo1-/+ mice in middle-age and old-age. The analysis indicated activations of kinase signaling p42/44 MAPK ERK1/2, AMPK, JNK, Wnt signaling via GSK3 inactivation, as well as increases of p-TAU and other AD biomarkers, PLCG1, EGFR, MET, Neurofibromin and RAS. Immune activation markers CD45 and CD31 were also elevated in the microenvironment. A majority of activated growth signaling components are of the oncogenic MET signaling complex. The discovery supports repurposing of cancer drugs targeting the MET signaling complex and EGFR-RAS-MAPK axis for intervention and/or treatment of genomic instability-driven AD.

## 1. Introduction

Five percent of Alzheimer’s disease (AD) develops early in life (familial/early-onset AD [EOAD]), which are facilitated by mutations in Amyloid Precursor Protein (APP), Presenilin 1, Tau, or APOE genes. The remaining 95% of human AD is late onset (LOAD) [1]. Although frequently-mutated genes are being identified and the effects on LOAD development are being assessed (e.g., [2,3]), the exact cause of LOAD remains unclear, and LOAD is considered multifactorial. With the lack of mechanistic understanding on the development, the intervention drug for LOAD has not been available in clinic. Yet, due to highly consistent presence of the major pathologies of amyloid plaques and neurofibrillary tangles in both EOAD and LOAD patients, drug-mediated removal of the pathologies has been a main goal for curative treatment of AD. As of early 2023, recent approvals of amyloid-targeting drugs for clinical use by US FDA provide hope for curative treatment of AD (e.g., Aducanumab; [4]. Lecanemab; [5]). However, because the amyloid-targeting AD drugs are still new in clinic, issues such as long-term efficacy, cost-effectiveness, or insurance coverage and drug availability are to be determined [6]. Hence, elucidation of LOAD development mechanism, and alternative strategies for AD intervention and treatment, are still unmet needs.

Genomic instability was identified as one of the twelve updated hallmarks of aging [7]. Genomic instability is an indicator of aging, as genomic instability or resulting aneuploidy is observed in a higher degree with aging [8], and also a driver of aging, as genomic instability models can be progeric (e.g., [9]). As human LOAD and Mild Cognitive Impairment (MCI) patients indicate a high degree of genomic instability in the brain and in lymphocytes (e.g., [10, 11]), we hypothesized that genomic instability facilitates LOAD pathology development, and discovered that a genomic instability mouse model (cohesinopathy-mediated Chromosome Instability [CIN] model Sgo1-/+) accumulated amyloid-beta (Abeta) in the brain beginning in late middle age (15-18 months of age) [12,13]. Thus, cohesinopathy-mediated genomic instability is suggested to trigger Abeta accumulation and/or drive LOAD. Additional studies revealed other similarities in pathological and molecular traits between human AD and the aged Sgo1-/+ mouse model [14]. It should be noted that Sgo1-/+ mice have shown LOAD pathology even with unaltered murine App (mApp). Due to three amino acid difference, mApp has a weaker ability to support Abeta generation or aggregation than human(ized) App (hApp), which is causing additional difficulty in modeling AD in mice [15]. It is plausible that the effects of Sgo1-/+-associated genomic instability will be demonstrated with hApp in a fuller manner in the mouse model.

Based on the transgenic mice and available reports on human AD, we built a hypothesis “the Amyloid-beta accumulation cycle” to explain how amyloid-beta accumulation occurs with genomic instability [16]. The hypothesis is built on following facts; (a) although the brain is generally considered a mitotically static organ with the exception of limited stem cell divisions, presence of mitotic signatures is observed in AD brains. (b) Amyloid-beta is cytotoxic, stress response-activating, and aneuploidgenic. (c) In cycling cells, amyloid-beta is accumulated in the mitotic phase. (d) Amyloid-beta possesses prionlike self-aggregation ability. The hypothesis purports that; these traits of amyloid can create autonomous mitotic cycle leading to “mitogenic signaling activation, mitotic cycle entry, aneuploidgenic mitotic interference by external and internal amyloid, mitotic catastrophe, amyloid release to microenvironment, and another round of mitotic cycle activation”, eventually leading to stromal accumulation of amyloid itself (presented in Fig1 A “the Amyloid-beta accumulation cycle”). When we observed that amyloid accumulation is only in Sgo1-/+, but not in BubR1-/+, we reasoned that Sgo1-/+ with intact spindle checkpoint is more prone to prolonged mitosis and mitotic catastrophe, compared with BubR1-/+ with defective spindle checkpoint and tendency to mitotic escape [13,16]. Thus, mitosis-prolonging type of genetic predisposition, such as cohesinopathy, would be more triggering to the amyloid-beta accumulation cycle.

**Figure 1.**
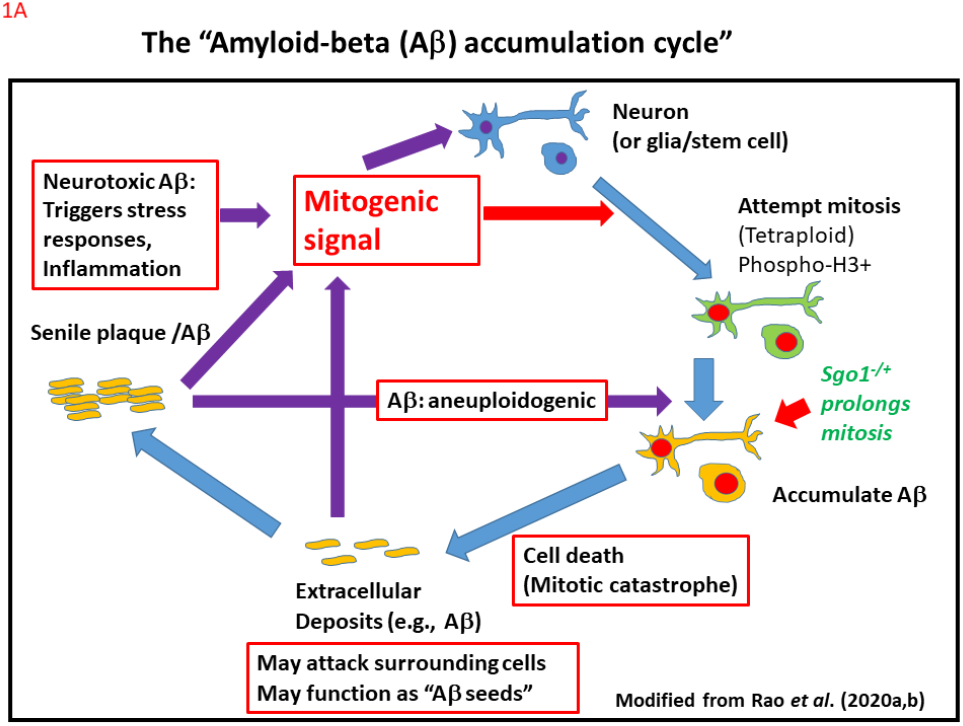
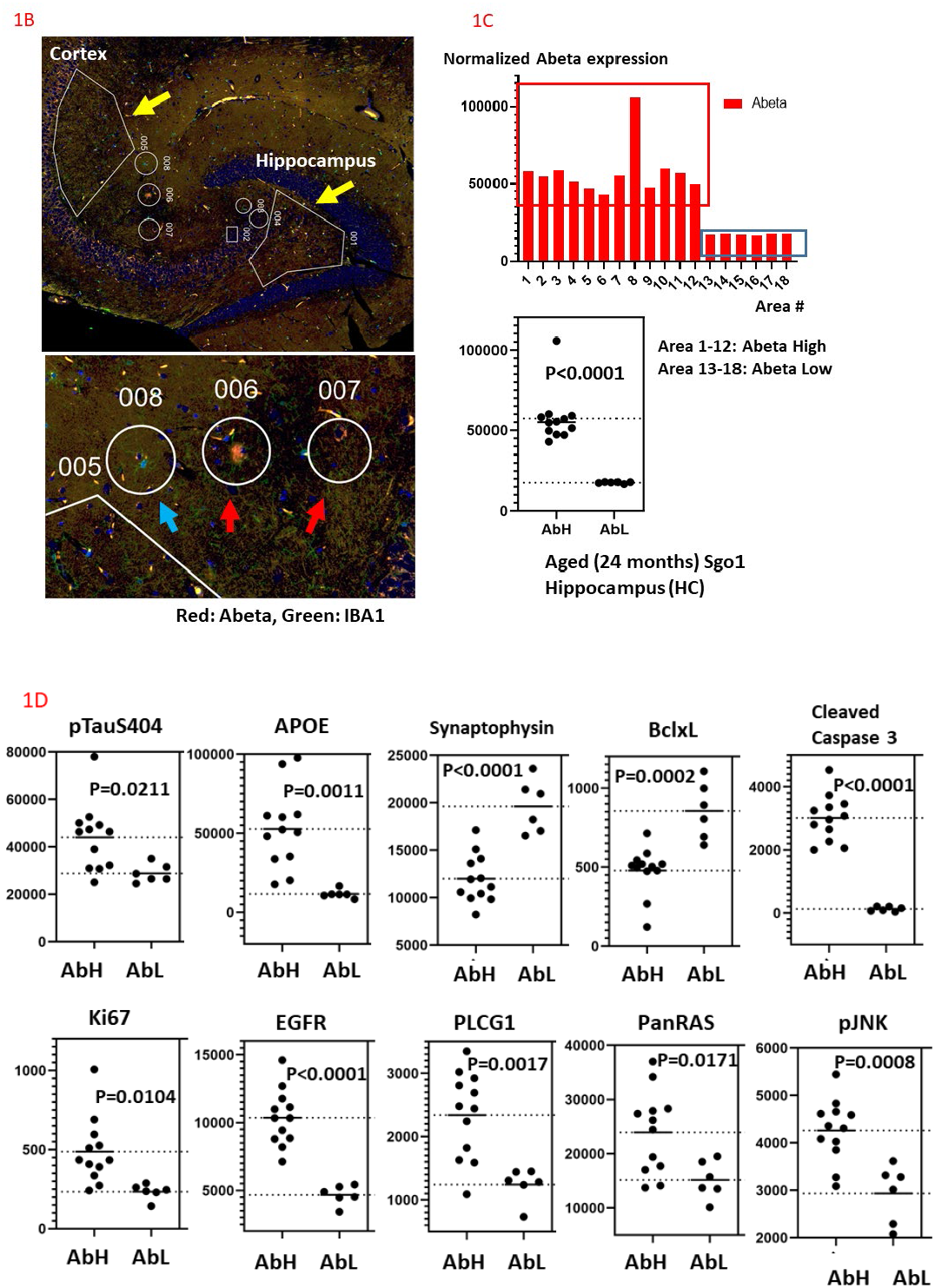
Areas in the vicinity of amyloid indicated high expression of AD pathology, cell death, neurodegeneration, and activation of MET growth signaling components. **(A) The “Amyloid-beta accumulation cycle”** The hypothesis was proposed in Rao et al., (2020a) [14] and GSK3 inactivation was identified as a candidate trigger of the cycle in the middle age (Rao et al., 2020b) [16]. However, the nature of mitogenic signaling was unidentified. **(B) Spatial analysis focused on amyloid and IBA1** Typical example of aged Sgo1-/+ brain used for the spatial analysis. Large ROI (yellow arrow) and small ROI (blue arrow focusing on IBA1, red arrow focusing on amyloid) were selected from amyloid-expressing Hippocampus and Cortex for quantification of proteins of interest. **(C) Comparison between Amyloid-beta-High (AbH) area and Amyloid-beta-Low (AbL) area** Based on expression level of amyloid, small ROI were categorized to AbH area or AbL area. Collectively, the amyloid expression levels were significantly different (p<0.0001). **(D)** Notable changes indicative of AbH microenvironment Expression of proteins of interest in AbH and AbL areas were compared. Presented here, are the results from aged Sgo1-/+ Hippocampus; increases in AD biomarkers (pTAU-S404, APOE), decrease in synapse marker Synaptophysin, decrease in anti-apoptotic Bclxl, increase in cell death marker cleaved Caspase 3, increase in growth marker Ki67, and increases in MET signaling components (EGFR, PLCG1, RAS) and its downstream (pJNK) (also see Table 2 and Table 3 for other components).

The amyloid-beta accumulation cycle hypothesis is an integrated hypothesis, and can accommodate many other factors and observations, such as ROS and other challenges to trigger damage and stress response, age-associated decrease in amyloid clearance, and higher presence of aneuploid cells in AD brains.

A key event in the “Amyloid-beta accumulation cycle” hypothesis is activation of mitogenic growth signaling. Yet, in the hypothesis, the nature of the “mitogenic signaling” was poorly described. In our previous report [14], we identified GSK3 inactivation in the middle age as a culprit of the trigger for amyloid-beta accumulation cycle. GSK3 inactivation can activate canonical Wnt signaling and can also cause accumulation of Activity-regulated cytoskeleton-associated protein (ARC) [17] (Wu et al., 2011), both of which would facilitate amyloid-beta increase. However, GSK3 is considered as a driver of later stage AD pathology (e.g., Tau phosphorylation) [18] and whether the inactivation of GSK3 is limited to middle age or not was unclear.

In this study, we focused our analysis on growth signaling proteins in the vicinity of amyloid-positive cells in aged Sgo1-/+ brains, using digital spatial profiling technology, and identified growth signaling that activated nearby amyloid-positive cells.

## 2. Materials and Methods

### Mice

C57/BL6-based Sgo1-/+ and control littermate wildtype (Sgo1+/+) mice were maintained for 18 or 24 months without any experimental treatment in the OUHSC rodent barrier facility, where clean and temperature/circadian cycle-controlled environment is maintained. Regular diet was used without modification and filtered water was available ad libitum. At the end of maintenance, the mice were euthanized with CO2 and brain samples were collected, fixed with 10% buffered formalin and embedded in paraffin to generate FFPE tissue samples at the OUHSC CCPDD histology core. The FFPE cassettes were submitted to the OMRF imaging core and mounted to the slides for NanoString DSP analysis.

### Brain tissue samples

Brain samples from age (24 or 18 months of age) and gender-matched (all female) groups were compared. Female samples were used, as human AD cases are more frequent in aged female than in aged males, although in our model such gender difference was not apparent. Group 1 (24 months of age, female Sgo1-/+, n=3), Group 2 (24 months of age, female wild type, n=3), Group 3 (18 months of age, female Sgo1-/+, n=4), Group 4 (18 months of age, female wild type, n=2). From each brain, Hippocampus (HC) and rear Cortex (C) were analyzed. For each HC or C, two large Region of interest (ROI), covering a majority of HC or comparable size of Cortex, and six small ROI, each covering an area of nearby morphology markers of amyloid or IBA1, were selected. As this analysis was focusing on molecular events nearby amyloid-beta pathology, main interest was on aged transgenic animals with amyloid accumulation, rather than pathology-free animals (e.g., wild type, younger Sgo1-/+ animals less than 12 months of age).

### Protein panels

We used protein profiler analysis, using Neuroscience panel, Alzheimer’s disease panel, expanded AD panel, and Oncology panel (Supplementary Table S1). Specific antibodies used for the NanoString protein panel analysis are published on the NanoString company website (2022 version).

### NanoString DSP analysis

The GeoMX DSP system uses tissue sections labeled with proprietary UV-cleavable barcoded oligo-attached antibodies. Sections were immunostained with morphology/pathology markers, which guided ROI selection. After selection of region of interest (ROI)s, the ROIs were irradiated with UV, and released barcoded oligos were collected and sequenced. The oligo counts represented quantified amount of protein of interest after normalization. For morphology/pathology markers, we selected anti-amyloid-beta (1-42) (Amyloid pathology), anti-IBA1 (activated microglia), and DAPI (nuclei).

We followed manufacturer’s instructions. We used GeoMx DSP Manual Slide Preparation User Manual NAN-10150-01 (April 2022), GeoMx DSP NGS Readout User Manual (Library Preparation and GeoMx NGS Pipeline) MAN-10153-01 (April 2022), and QC and Normalization was performed with guidance from GeoMx DSP Data Analysis User Manual MAN-10154-01 for v2.5 software (March 2022). Optimal normalization standards were determined to be housekeeping gene proteins s6 and GAPDH, based on their strong correlation plot data.

### Sequencing Data analysis and statistical Analysis

The cleaved oligos were sequenced in the OMRF genomics core, analyzed with GeoMx DSP Data Analysis package, and normalized quantification data were presented in excel sheets format. We analyzed the normalized output and visualized them with GraphPad Prism software (ver9). Statistical significance was evaluated by algorithms integral to the aforementioned software with student’s t-test. FDR-adjusted P values of <0.05 were considered significant.

## 3. Results

### 3.1 Study design

To identify growth signaling involved in cell cycle progression in the brains of aged Sgo1-/+ genomic instability mice, we prepared the following samples; (a) Sgo1-/+ mouse brain 24 months (“aged”), Cortex and Hippocampus, (b) wild type (WT) mouse brain 24 months, Cortex and Hippocampus, (c) Sgo1-/+ mouse brain 18 months (“late middle age”), Cortex and Hippocampus, and (d) WT mouse brain 18 months, Cortex and Hippocampus. Then, we used NanoString GeoMX DSP protein profiler with Oncology panel, AD pathology panel, AD pathology extended panel, and neuroscience panel. The combined panels were designed to quantify 64 proteins of interest in specified areas (typical view shown in Fig1 B). From each group, we selected 18 small areas in the vicinity of amyloid-beta and/or of IBA1 as region of interest (ROIs) to test (i) whether specific signaling is/are activated in the vicinity of amyloid and (ii) whether amyloid and IBA1 localize close to each other, which would implicate microglia-mediated amyloid surveillance. Hence, we focused our analysis on signaling pathways nearby accumulating amyloid-beta, and immune surveillance on amyloid-beta-accumulating cells.

Excluding proteins for internal control purposes (e.g., Rabbit IgG2a, IgG2b, GAPDH, Histone H3), we categorized the 64 proteins in the panels to six functional groups (**Supplementary Table S1**); (i) AD-associated pathophysiological markers (15 proteins; i.e., Amyloid-beta 1-42, APP, p-TAU [S404, S199, S396, S214, S231], Tau, APOE, TMEM119, PSEN1, Neprilysin, IDE, BACE1, Tdp43), (ii) growth signaling (21 proteins including pJNK, p-GSK3A+B, p-p42/44 MAPK ERK1/2, p38MAPK, p-AMPK, p-MEK1, PLCG1, EGFR, BRAF, MET, neurofibromin, pan-RAS, Ki67, p-S6, p-PRAS40, p-AKT), (iii) neuronal function (9 proteins; GFAP, NeuN, Neurofilament L, P2FX7, synaptophysin, Myelin Basic Protein [MBP], MAP2, Olig2, NRGN), (iv) DNA damage (3 proteins; gammaH2AX, p21, p53), (v) cell death (6 including BAD, cleaved caspase 3, Bclxl, BAD, PARP, BIM), and (vi) immune function (3 proteins; CD45, CD31, IBA1).

In aged (24mo.) and late-middle-aged (18mo.) WT, amyloid-beta specific signals were poorly presented as predicted (not shown). Thus, we focused our analysis with AD pathology on aged (24mo.) and late-middle-aged (18mo.) Sgo1-/+ brain samples. Consistent with previous reports [13, 14], in the Sgo1-/+ brains, amyloid-specific signals were observed predominantly in the Hippocampus and rear Cortex. As such, ROIs were drawn in these areas.

### 3.2. Microenvironment in vicinity of amyloid-accumulating cells

As a major merit of a spatial analysis, signals can be distinguished with single cell resolution, allowing us to pinpoint areas of interest. Using amyloid expression, we identified high amyloid areas (amyloid-accumulating areas) and low amyloid areas (Fig.1C). Subsequently, we compared the amounts of other proteins of interest in high amyloid areas (i.e., “vicinity of amyloid” areas) and low amyloid areas.

#### Hippocampus

In aged Sgo1 brain Hippocampus, the following proteins were expressed higher in the vicinity of amyloid-accumulating cells (Fig.1D, Table 1); AD pathology proteins (p-Tau, Tau, BACE1, APOE, TMEM119, PSEN, Neprilysin, APP, IDE), growth signaling (p-JNK, p-GSK3, p-AMPK, PLCG1, EGFR, BRAF, MET, neurofibromin, RAS), Neurofilament L (neuronal damage marker), Cell death (BAD, cleaved Caspase3, BIM), Immune function (CD45, CD31).

**Table 1.**
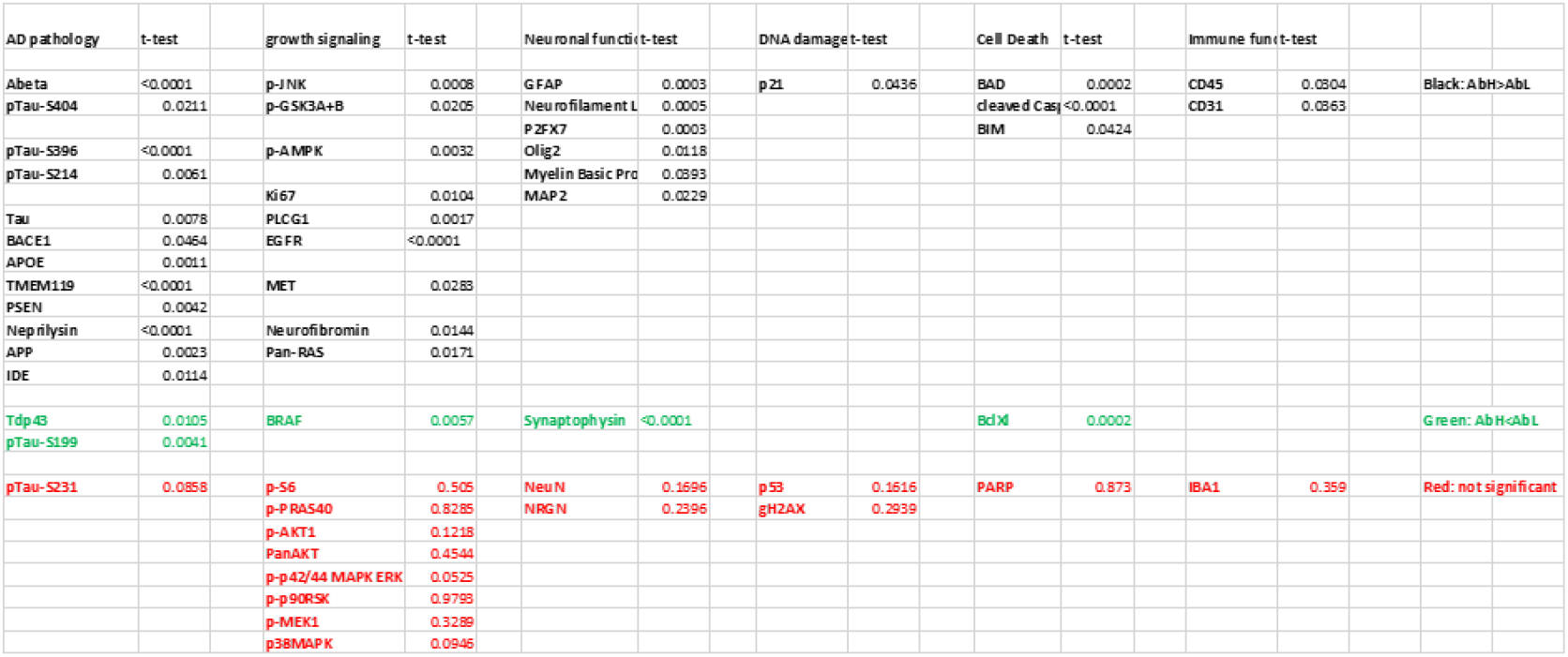
Amyloid-high vs Amyloid-low (Aged Sgo1 Hippocampus), comparisons of protein expression in Hippocampus.

In contrast, Synaptophysin (synapse marker), TDP43, BRAF, BclxL were expressed lower in the vicinity of amyloid-accumulating cells. Decrease in synaptophysin suggests loss of synapses. As TDP43 is involved in DNA double strand break repair, decrease in TDP43 may facilitate DNA damage accumulation. Decrease of anti-apoptotic BclxL may facilitate cell death.

The results indicate the following; (a) abnormal expressions and increases in AD pathology markers, (b) increased cell death with loss of anti-apoptotic BclxL and activation of Caspase 3, (c) loss of synaptophysin, specific marker for synaptic terminals, suggesting synapse loss, (d) increase in growth marker Ki67, suggesting abnormal mitotic cycle reentry, (e) modulated energy homeostasis and/or autophagy that may serve to improve survival (p-AMPK), and (f) increases in growth signaling components (e.g., EGFR, MET, PLCG1, RAS) and activation markers (p-JNK). Neurofibromin, a RAS-GAP and tumor suppressor also increases, suggesting RAS-inactivation is also activated. Yet, judging from activation of the downstream JNK and MAPK, effects of neurofibromin may be insufficient to negate growth signaling. Altogether, these changes in the vicinity of amyloid-accumulating cells point toward AD-pathogenic, cell death-facilitating, neurodegenerative and pro-mitogenic microenvironment.

#### Cortex

In aged Sgo1 brain Cortex, the following proteins were expressed higher in the vicinity of amyloid-accumulating cells (amyloid-high areas) (Table 2); AD pathology proteins (p-Tau, Tau, APOE, TMEM119, PSEN, Neprilysin, APP, IDE), growth signaling (p-JNK, p-GSK3A+B, p-p42/44 MAPK ERK, p-AMPK, p-MEK1, PLCG1, EGFR, MET, neurofibromin, RAS, Ki67), Neurofilament L (neuronal damage marker), Cell death (BAD, cleaved Caspase3), Immune function (CD45, CD31). There was significant overlapping with the profile in Hippocampus, with exceptions of p-p42/44 MAPK ERK1/2 and gamma H2AX being higher in Cortex only (Fig. 2A). In contrast, increases in proapoptotic BIM and senescence marker p21 were consistently specific to Hippocampus.

**Figure 2.**
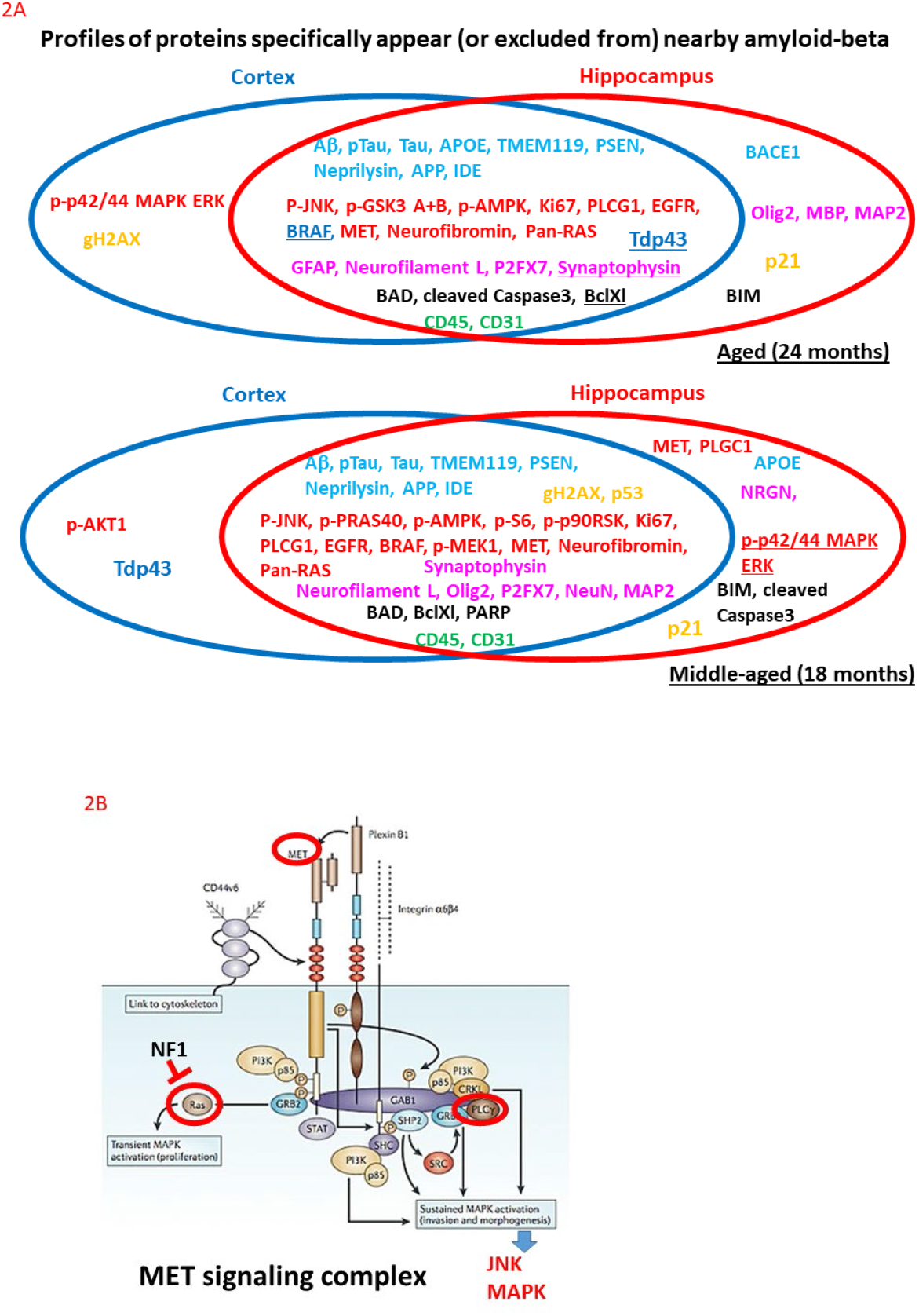
Identification of MET signaling as major mitogenic signaling. **(A) Profiles of proteins specifically appear (or excluded from) nearby amyloid-beta in aged and/or middle-aged Sgo1-/+ brains** In the Hippocampus and Cortex, there was a significant overlapping in the proteins specifically appear in the vicinity of amyloid. Four proteins were excluded from the vicinity of amyloid (underlined; BRAF, Synaptophysin, Tdp43, Bclxl). Common growth signaling components are identified both in aged and middle-aged animals (e.g., p-JNK, EGFR, MET, RAS), suggesting that they represent the key drivers of AD pathology. Signaling potentially antagonizing AD pathology (p-AMPK, Neurofibromin) are also present. Peculiarly, BRAF and Bclxl expression pattern changed toward opposite direction over time; in the middle age they are expressed higher around amyloid, but in old age they are expressed less. **(B) MET, EGFR, RAS, PLCG, JNK, MAPK are components of the MET signaling complex**.

### 3.3. Microenvironment in vicinity of IBA1-positive cells

As with amyloid-beta, we used IBA1 expression to identify IBA1-high area and IBA1low area.

We hypothesized that IBA1, a microglia activation marker, would co-localize to amyloid, suggesting that amyloid attracts and is targeted by active microglia as a part of immunosurveillance. However, in both Cortex and Hippocampus, high IBA1 was correlated with high CD45, a leukocyte marker (Supplementary Fig. S2A), but not with AD marker (amyloid, p-Tau) or any other proteins tested in aged Sgo1 Hippocampus (Supplementary Fig. S2B). Together, the results suggest that IBA1-positive activated microglia are not selectively localized in the vicinity of amyloid-accumulating cells.

Identified components (circled in red) are of the MET signaling complex (modified from Abounader et al., (2004) [19]). The MET complex is a multicomponent signaling complex, whose main hub is receptor Tyrosine Kinase MET. MET interacts with other components including GAB1, SRC, SHC, RAS, PLCG, PI3K, STAT, and transduce signal to the downstream JNK, MAPK. The MET complex can crosstalk with EGFR (e.g., [20]). Under normal circumstances, the MET complex is involved in embryonic development, organogenesis and wound healing, while abnormally activated MET triggers tumor growth, angiogenesis and metastasis. NF1 (Neurofibromin) is a RAS-GAP that inhibits RAS activation [21].

AD pathology proteins (p-Tau, Tau, BACE1, APOE, TMEM119, PSEN, Neprilysin, APP, IDE), growth signaling (p-JNK, p-GSK3, p-AMPK, PLCG1, EGFR, BRAF, MET, neurofibromin, RAS), Neurofilament L (neuronal damage marker), Cell death (BAD, cleaved Caspase3, BIM), Immune function (CD45, CD31) were abnormally expressed in the vicinity of amyloid-accumulating microenvironment at higher levels. Synaptophysin (synapse marker), TDP43, BRAF and BclXL were expressed at lower levels. Black: Higher expression in Amyloid-high areas (p<0.05). Green: Lower expression in Amyloid-high areas. Red: expression level difference is not significant.

Table 2. Table 2 Amyloid-high vs Amyloid-low (Aged Sgo1 Cortex), comparisons of protein expression in rear Cortex.

AD pathology proteins (p-Tau, Tau, APOE, TMEM119, PSEN, Neprilysin, APP, IDE), growth signaling (p-JNK, p-GSK3, p-p42/44 MAPK ERK, p-AMPK, p-MEK1, PLCG1, EGFR, MET, neurofibromin, RAS, Ki67), Neurofilament L (neuronal damage marker), Cell death (BAD, cleaved Caspase3), Immune function (CD45, CD31) were abnormally expressed in the vicinity of amyloid-accumulating microenvironment at higher levels. Synaptophysin (synapse marker), TDP43, BRAF and BclXL were expressed at lower levels. Black: Higher expression in Amyloid-high areas. Green: Lower expression in Amyloid-high areas. Red: expression level difference is not significant.

## 4. Discussion

In recent years, spatial analysis focusing on microenvironment has secured the position of an invaluable tool to analyze pathogenic events at a higher resolution. We applied the technique to transgenic cohesinopathy mice brains that are accumulating amyloid, so that we can obtain insight in genomic instability-associated AD pathology driving events. Prevalence of aneuploidy in human AD brains [10,11] implicate relevance and applicability of results of this study to human AD pathology development. Unlike transgenic early-onset AD model mice, which are APP-focused models, the cohesinopathy mice do not overproduce APP nor carry knock-in mutations of EOAD in APP or APOE. Thus, we anticipate that the Sgo1-/+ mice may represent some aspects of human sporadic late-onset AD, which also do not frequently carry mutations in APP (e.g., [22]).

We hypothesized that growth signaling is activated in amyloid-accumulating microenvironment and driving pathogenic mitotic cycles. This study focused on AD pathology occurring in aged Sgo1-/+ mice brains, and confirmed activation of growth signaling components, namely, phosphorylation-mediated activations of JNK, MAPK, and AMPK, inactivation of GSK3A+B (which activates canonical Wnt signaling), and increases in PLCG1 (phospholipase C, gamma 1), EGFR (Epidermal growth factor receptor), MET (proto-oncogene, receptor TK), Neurofibromin (RAS-GAP, tumor suppressor regulating RAS/MAPK), and RAS.

Notably, these components form a large receptor tyrosine kinase complex, the MET signaling complex, or are a part of its cross-talking signaling (e.g., EGFR). The MET complex activates its downstream JNK-MAPK signaling (Fig. 2B). As other tested signaling components did not show signs of activation (e.g., p-S6, p-PRAS40, p-AKT, p-p90RSK, pMEK1), clear distinction among the tested signaling pathways was observed.

Human AD brains show signs of mitotic progression as a part of pathology [23]. However, the mitotic progression has not been a mainstream target for AD drug development. With the Amyloid-beta accumulation cycle hypothesis, we proposed to target the cycle. And an option to target the cycle was the use of CDK inhibitors [24]. Although CDK inhibitors indicated amelioration effects on mice AD models [25, 26] and strong association with AD clinical and pathological traits and cell cycle-enriched module/subnetwork was identified with human AD gene expression analysis [27], to our knowledge clinical trials of CDK inhibitors on AD have not occurred or the results reported, as of early 2023. Yet, as amyloid-targeting drugs are new to the clinic and have much to prove in clinical settings, another class of AD drug is still in need. In this study, we identified another component of the amyloid-beta accumulation cycle, which are the MET signaling complex [19, 28], RAS, EGFR, PLCG and its downstream, the JNK-MAPK signaling. Hence, we propose that the MET complex, RAS, EGFR, PLCG and the JNK-MAPK signaling are new targets to inhibit the amyloid-accumulation cycle and AD intervention drug targets.

Indeed, recent studies suggest that the MET complex and its components may be involved in human AD. MET and its ligand HGF are involved in synaptogenesis, neuronal growth and survival, and deregulation of MET is associated with developmental disorders such as autism spectrum disorder [29], supporting the notion that the MET complex is a signaling utilized in damage response and tissue maintenance under normal circumstances in adult brains. The HGF/c-MET axis was proposed as a potential AD drug target, although in the context of synaptogenesis regulation [30].

PLCG1 is known to be associated with AD pathogenesis [31]. Human AD-specific single-nucleotide variants and abnormal exon splicing of the PLCG1 gene were discovered through deep learning-based prediction on GWAS database [32]. Although the mechanism was thought to be through regulating neuronal functions, involvements of PLCG1 on AD pathogenesis may be open to new interpretation, through the viewpoint of growth regulation. In addition, RAS-MAPK signaling components were identified as DEGs in human AD gene expression data mining [33]. Zhang et al., (2021) [34] reported that downregulation of EIF3H, RAD51C, FAM162A, BLVRA, ATP6V1H, and BRAF was closely related to the occurrence of human AD, corroborating with our results of BRAF downregulation.

Purported herbal AD medicine can target EGFR and other growth signaling. For example, ginseng is medicinal herb claimed to work for age-associated diseases including AD. A total of 22 bioactive compounds were identified from ginseng, and their molecular targets include growth signaling (e.g., Ras signaling, PI3K-AKT signaling). The compound-target-GO-route network found EGFR, MAPK1, MAPK14, AKT1, CASP3, and PRKACA as key genes, with PI3K-AKT signaling being the most important pathway for ginseng’s anti-AD activity [35]. Synsepalum dulcificum, also known as the miracle fruit, is a hopeful of herbal AD treatment. The seed extract indicated ameliorating effects on learning memory functions of 2xFAD AD model mice. The molecular docking predictions showed that the seed compounds were stably bound to core targets, specifically AKT1, EGFR, ESR1, PPARA, and PPARG. Thus, the seeds are discussed to potentially play a therapeutic role in AD by affecting the insulin signaling pathway and the Wnt pathway [36]. Ginkgo biloba L. leaves have shown potentially beneficial effects for people with dementia, when it is administered at doses greater than 200mg/day for at least 5 months [37]. Twenty-seven active components of Ginkgo biloba leaves were implicated to affect 120 metabolic pathways, such as the PI3K/AKT pathway, by regulating 147 targets, including AKT1, ALB, HSP90AA1, PTGS2, MMP9, APP and EGFR [38]. Genistein is an isoflavone from soybeans, and is an inhibitor of EGFR. Genistein supplementation was effective to treat the APP/PS1 AD mouse model, and in a clinical trial, genistein treatment resulted in a significant improvement in two of the tests used and a tendency to improve in all the rest of them. The amyloid-beta deposition analysis showed that genistein-treated patients did not increase their uptake in the anterior cingulate gyrus after treatment, while placebo-treated did increase it [39]. Thus, there emerged the common target EGFR.

Small molecule EGFR inhibitors (e.g., erlotinib (Tarceva), gefitinib (Iressa), afatinib (Gilotrif), dacomitinib (Vizimpro), and osimertinib (Tagrisso)) are clinically used as chemotherapy drugs for cancers including NSCLC (reviewed in Zubair & Bandyopadhyay, 2023 [40]). The drugs are candidates for repurposing. While blood-brain barrier (BBB) impairment is widely reported in AD patients [41], drug delivery to the brain is an issue to consider. In mouse and rhesus macaques, co-infusion treatment of erlotinib with Tariquidar, a P-glycoprotein inhibitor, allowed erlotinib to pass through BBB and accumulation in the brain [42], hence showing a promise in solving drug delivery issue toward feasible clinical application.

In addition to the MET growth signaling, this study showed AMPK activation and increase of neurofibromin in amyloid-high areas. Activation of AMPK is a sign of autophagy pathway activation, which can antagonize amyloid toxicity [43]. Neurofibromin is a RAS-GAP, antagonizing RAS signaling activation, thus function as tumor suppressor [21]. They may also represent pathways that can be augmented to put a break on the amyloidbeta accumulation cycle.

Overall, this study indicated activation of the MET signaling complex, EGFR, and downstream RAS, PLCG, JNK/MAPK in the vicinity of amyloid-beta accumulating cells, thus providing supporting evidence for the “Amyloid-beta accumulation cycle” hypothesis. At the same time, the results suggest the MET complex and EGFR, RAS, PLCG and JNK/MAPK signaling as drug targets to shut down the cycle, in addition to amyloid itself and cell cycle-driving CDKs. The MET complex is a signaling hub and a convergence point of growth signaling at the cell membrane. As such, a number of targeting drugs have been developed already and in clinical use. We propose that repurposing of such drugs (e.g., EGFR inhibitor) to target the MET signaling complex and the downstream pathway (RAS, PLCG1, JNK, MAPK) will serve as intervention of or adjuvant treatment for LOAD. This unexplored approach may lead to innovation in clinical practice if proven.

As limitation, we have not been able to determine whether the MET signaling components, such as RAS or EGFR, have acquired activating mutation(s) in or in the vicinity of amyloid-accumulating microenvironment, as in the case with various cancers [44, 45]. Hence, whether drugs specific for activating mutation(s) (e.g., EGFR-L858R, KRAS-G12C or G12D) would indicate efficacy or not remains to be determined. Another limitation is that the protein panel is not exhaustive and for select proteins only. Thus, from this assay, activation of other predicted components of the Amyloid-beta accumulation cycle (e.g., CDKs) were not determined. Also, the effects of Sgo1 haploinsufficiency has been analyzed only in C57BL/6 background (inbred, non-tumor-prone) thus far, and other strain background, which may influence phenotype, have not been tested.

## Supporting information

Supplementary Table S2 Sgo18HC

Supplementary Table S3 Sgo18C

Table 1 Sgo24HC

Table 2 Sgo24C

## Supplementary Materials

The following supporting information can be downloaded at: xxxx **Supplementary Figure S1:** Selection of Amyloid-High area and Amyloid-Low area in middle aged samples. Amyloid-high area and Amyloid-low areas were determined based on amyloid expression. The expression levels are significantly different (P<0.0001). (A) Middle-aged (18 months) Sgo1 brain Hippocampus. (B) Middle-aged (18 months) Sgo1 brain Cortex. **Supplementary Figure S2:** IBA1 high and low areas were identified and amount of proteins of interest were compared. (A) IBA1 high area expressed higher amount of CD45. (B) IBA1 expression levels did not correlate with amyloid-beta or phospho-TAU (S404). **Supplementary Table S1:** List of 64 proteins quantified in this study. **Supplementary Table S2:** Amyloid-high vs Amyloid-low (middle-Aged Sgo1 Hippocampus), comparisons of protein expression in Hippocampus. AD pathology proteins (p-Tau, Tau, TMEM119, PSEN, APOE, APP, IDE, Neprilysin), growth signaling (p-JNK,p-AKT1, p-PRAS40, pAMPK, p-S6, p-p90RSK, EGFR, BRAF, p-MEK1, neurofibromin, RAS, AKT, p38MAPK, MET, Ki67), Neurofilament L (neuronal damage marker), Cell death (BAD, BIM, BclXl, PARP, cleaved Caspase 3), Immune function (CD45, CD31) were expressed higher in amyloid-accumulating microenvironment. While, p-P42/44 MAPK ERK and MAP2 were expressed lower. **Supplementary Table S3:** Amyloid-high vs Amyloid-low (middle-Aged Sgo1 Cortex), comparisons of protein expression in Cortex. AD pathology proteins (p-Tau, Tau, TMEM119, PSEN, APP, IDE, Neprilysin), growth signaling (p-JNK, p-AKT1, p-PRAS40, p-AMPK, p-S6, p-p90RSK, p-P42/44 MAPK ERK, EGFR, BRAF, p-MEK1, neurofibromin, RAS, AKT, p38MAPK), Neurofilament L (neuronal damage marker), Cell death (BAD, BIM, BclXl, PARP), Immune function (CD45, CD31) were expressed higher in amyloid-accumulating microenvironment. Synaptophysin (synapse marker) and TDP43 were expressed lower in amyloid-high areas in middle-age Cortex.

## Author Contributions

Conceptualization, H.Y.Y.; methodology, H.Y.Y., J.C., B.F.; software, J. C., B. F.; validation, H.Y.Y., J.C., B.F.; formal analysis, H.Y.Y., J.C., B.F.; investigation, H.Y.Y., J.C., B.F. Y.Z.; resources, C.V.R.,Y.Z., J.C., B.F., H.Y.Y.; data curation, J.C., B.F.; writing—original draft preparation, H.Y.Y.; writing—review and editing, H.Y.Y., C.V.R., J.C., B.F.; visualization, H.Y.Y., J.C., B.F.; supervision, H.Y.Y.; project administration, H.Y.Y., C.V.R.; funding acquisition, H.Y.Y., C.V.R. All authors have read and agreed to the published version of the manuscript.

## Funding

This research was funded by the Presbyterian Health Foundation (PHF) bridge grant 2021. The APC was funded by XXX/TBD.

## Institutional Review Board Statement

Mice were maintained and handled following the OUHSC Institutional Animal Care and Use Committee (IACUC) guidelines that conform to national guidelines for animal usage in research. The animal study protocol was approved by the IACUC of OUHSC (Protocol #: 21-066, approved Sept 22, 2021).

## Informed Consent Statement

Not Applicable. No human subject was used.

## Data Availability Statement

The sequencing datasets from NanoString analysis will be available from the corresponding author by a reasonable request. The reagents and animals described in this article are available under a material transfer agreement with University of Oklahoma Health Sciences Center.

## Acknowledgments

In this section, you can acknowledge any support given which is not covered by the author contribution or funding sections. This may include administrative and technical support, or donations in kind (e.g., materials used for experiments).

## Conflicts of Interest

The authors declare no conflict of interest. The funders had no role in the design of the study; in the collection, analyses, or interpretation of data; in the writing of the manuscript; or in the decision to publish the results.

## Disclaimer/Publisher’s Note

The statements, opinions and data contained in all publications are solely those of the individual author(s) and contributor(s) and not of xxxx and/or the editor(s). xxxx and/or the editor(s) disclaim responsibility for any injury to people or property resulting from any ideas, methods, instructions or products referred to in the content.

## Notes

### Competing Interest Statement

The authors have declared no competing interest.

## References

1. Scheltens, P., Blennow, K., Breteler, M.M., de Strooper, B., Frisoni, G.B., Salloway, S, & Van der Flier, W.M. (2016). Alzheimer’s disease. Lancet, 388, 505–17.

2. Andrade-Guerrero, J., Santiago-Balmaseda, A., Jeronimo-Aguilar, P., Vargas-Rodríguez, I., Cadena-Suárez, A.R., Sánchez-Garibay, C., Pozo-Molina, G., Méndez-Catalá, C.F., Cardenas-Aguayo, M.D., Diaz-Cintra, S., Pacheco-Herrero, M., Luna-Muñoz, J., & Soto-Rojas, L.O. (2023). Alzheimer’s Disease: An Updated Overview of Its Genetics. Int J Mol Sci, 24, 3754.

3. Oblak, A.L., Forner, S., Territo, P.R., Sasner, M., Carter, G.W., Howell, G.R., Sukoff-Rizzo, S.J., Logsdon, B.A., Mangravite, L.M., Mortazavi, A., Baglietto-Vargas, D., Green, K.N., MacGregor, G.R., Wood, M.A., Tenner, A.J., LaFerla, F.M., & Lamb, B.T.; and The MODEL-AD Consortium. (2020). Model organism development and evaluation for late-onset Alzheimer’s disease: MODEL-AD. Alzheimers Dement (N Y), 6, e12110.

4. Rahman, A., Hossen, M.A., Chowdhury, M.F.I., Bari, S., Tamanna, N., Sultana, S.S., Haque, S.N., & Saif-Ur-Rahman, K.M. (2023). Aducanumab for the treatment of Alzheimer’s disease: a systematic review. Psychogeriatrics, Feb 12. doi: 10.1111/psyg.12944. Online ahead of print. PMID: 36775284

5. van Dyck, C.H., Swanson, C.J., Aisen, P., Bateman, R.J., Chen, C., Gee, M., Kanekiyo, M., Li, D., Reyderman, L., Cohen, S., Froelich, L., Katayama, S., Sabbagh, M., Vellas, B., Watson, D., Dhadda, S., Irizarry, M., Kramer, L.D., & Iwatsubo, T. (2023). Lecanemab in Early Alzheimer’s Disease. N Engl J Med, 388, 9–21.

6. Lin, P.J., Levine, A., Rucker, J., & Chambers, J.D. (2023). Variation in Medicaid and commercial payer coverage of aducanumab for Alzheimer’s disease. Alzheimers Dement. Feb 28. doi: 10.1002/alz.12965. Online ahead of print.

7. López-Otín, C., Blasco, M.A., Partridge, L., Serrano, M., & Kroemer, G. (2023). Hallmarks of aging: An expanding universe. Cell, 186, 243–278.

8. Tabula Muris Consortium. (2020). A single-cell transcriptomic atlas characterizes ageing tissues in the mouse. Nature, 583, 590–595.

9. Baker, D.J., Perez-Terzic, C., Jin, F., Pitel, K.S., Niederländer, N.J., Jeganathan, K., Yamada, S., Reyes, S., Rowe, L., Hiddinga, H.J., Eberhardt, N.L., Terzic, A., & van Deursen, J.M. (2008). Opposing roles for p16Ink4a and p19Arf in senescence and ageing caused by BubR1 insufficiency. Nat Cell Biol, 10, 825–36.

10. Bajic, V., Spremo-Potparevic, B., Zivkovic, L., Isenovic, E.R., & Arendt, T. (2015). Cohesion and the aneuploid phenotype in Alzheimer’s disease: A tale of genome instability. Neuroscience and Biobehavioral Reviews, 55, 365–374.

11. Hou, Y., Song, H., Croteau, D. L., Akbari, M., & Bohr, V. A. (2017). Genome instability in Alzheimer disease. Mechanisms of Ageing and Development, 161, 83–94.

12. Rao, C.V., Farooqui, M., Zhang, Y., Asch, A.S., & Yamada, H.Y. (2018a). Spontaneous development of Alzheimer’s disease-associated brain pathology in a Shugoshin-1 mouse cohesinopathy model. Aging Cell, 17, e12797.

13. Rao, C.V., Farooqui, M., Asch, A.S., & Yamada, H.Y. (2018b). Critical role of mitosis in spontaneous late-onset Alzheimer’s disease; from a Shugoshin 1 cohesinopathy mouse model. Cell Cycle, 17, 2321–2334.

14. Rao, C.V., Farooqui, M., Madhavaram, A., Zhang, Y., Asch, A.S., & Yamada, H.Y. (2020a). GSK3-ARC/Arg3.1 and GSK3-Wnt signaling axes trigger amyloid-β accumulation and neuroinflammation in middle-aged Shugoshin 1 mice. Aging Cell, 19, e13221.

15. Ueno, H., Yamaguchi, T., Fukunaga, S., Okada, Y., Yano, Y., Hoshino, M., & Matsuzaki, K. (2014). Comparison between the aggregation of human and rodent amyloid beta-proteins in GM1 ganglioside clusters. Biochemistry, 53, 7523–30.

16. Rao, C.V., Asch, A.S., Carr, D.J.J., & Yamada, H.Y. (2020b). “Amyloid-beta accumulation cycle” as a prevention and/or therapy target for Alzheimer’s disease. Aging Cell, 19, e13109.

17. Wu, J., Petralia, R.S., Kurushima, H., Patel, H., Jung, M.Y., Volk, L., Chowdhury, S., Shepherd, J.D., Dehoff, M., Li, Y., Kuhl, D., Huganir, R.L., Price, D.L., Scannevin, R., Troncoso, J.C., Wong, P.C., & Worley, P.F. (2011). Arc/Arg3.1 regulates an endosomal pathway essential for activity-dependent beta-amyloid generation. Cell, 147, 615–28.

18. Lauretti, E., Dincer, O., & Praticò, D. (2020). Glycogen synthase kinase-3 signaling in Alzheimer’s disease. Biochim Biophys Acta Mol Cell Res, 1867, 118664.

19. Abounader, R., Reznik, T., Colantuoni, C., Martinez-Murillo, F., Rosen, E.M., & Laterra, J. (2004). Regulation of c-Met-dependent gene expression by PTEN. Oncogene, 23, 9173–82.

20. Meng, X., Zhao, Y., Han, B., Zha, C., Zhang, Y., Li, Z., Wu, P., Qi, T., Jiang, C., Liu, Y., & Cai, J. (2020). Dual functionalized brain-targeting nanoinhibitors restrain temozolomide-resistant glioma via attenuating EGFR and MET signaling pathways. Nat Commun, 11, 594.

21. Mo, J., Moye, S.L., McKay, R.M., & Le, L.Q. (2022). Neurofibromin and suppression of tumorigenesis: beyond the GAP. Oncogene, 41, 1235–1251.

22. Arango, D., Cruts, M., Torres, O., Backhovens, H., Serrano, M.L., Villareal, E., Montañes, P., Matallana, D., Cano, C., Van Broeckhoven, C., & Jacquier, M. (2001). Systematic genetic study of Alzheimer disease in Latin America: mutation frequencies of the amyloid beta precursor protein and presenilin genes in Colombia. Am J Med Genet, 103, 138–43.

23. Herrup, K. (2010). The involvement of cell cycle events in the pathogenesis of Alzheimer’s disease. Alzheimer’s Research & Therapy, 2, 13.

24. Panagiotou, E., Gomatou, G., Trontzas, I.P., Syrigos, N., & Kotteas, E. (2022). Cyclin-dependent kinase (CDK) inhibitors in solid tumors: a review of clinical trials. Clin Transl Oncol, 24, 161–192.

25. Leggio, G. M., Catania, M. V., Puzzo, D., Spatuzza, M., Pellitteri, R., Gulisano, W., Torrisi, S.A., Giurdanella, G., Piazza, C., Impellizzeri, A.R., Gozzo, L., Navarria, A., Bucolo, C., Nicoletti, F., Palmeri, A., Salomone, S., Copani, A., Caraci, F., & Drago, F. (2016). The antineoplastic drug flavopiridol reverses memory impairment induced by Amyloid-ß1-42 oligomers in mice. Pharmacological Research, 106, 10–20.

26. Huang, F., Wang, M., Liu, R., Wang, J.-Z., Schadt, E., Haroutunian, V., Katsel, P., Zhang, B., & Wang, X. (2019). CDT2-controlled cell cycle reentry regulates the pathogenesis of Alzheimer’s disease. Alzheimer’s & Dementia: The Journal of the Alzheimer’s Association, 15, 217–231.

27. Zhang, B., Gaiteri, C., Bodea, L.-G., Wang, Z., McElwee, J., Podtelezhnikov, A. A., Zhang, C., Xie, T., Tran, L., Dobrin, R., Fluder, E., Clurman, B., Melquist, S., Narayanan, M., Suver, C., Shah, H., Mahajan, M., Gillis, T., Mysore, J., MacDonald, M.E., Lamb, J.R., Bennett, D.A., Molony, C., Stone, D.J., Gudnason, V., Myers, A.J., Schadt, E.E., Neumann, H., Zhu, J., & Emilsson, V. (2013). Integrated systems approach identifies genetic nodes and networks in late-onset Alzheimer’s disease. Cell, 153, 707– 720.

28. Yao, H.P., Tong, X.M., & Wang, M.H. (2021). Oncogenic mechanism-based pharmaceutical validation of therapeutics targeting MET receptor tyrosine kinase. Ther Adv Med Oncol, 13, 17588359211006957.

29. Desole, C., Gallo, S., Vitacolonna, A., Montarolo, F., Bertolotto, A., Vivien, D., Comoglio, P., & Crepaldi, T. (2021). HGF and MET: From Brain Development to Neurological Disorders. Front Cell Dev Biol, 9, 683609.

30. Wright, J.W., & Harding, J.W. (2015). The Brain Hepatocyte Growth Factor/c-Met Receptor System: A New Target for the Treatment of Alzheimer’s Disease. J Alzheimers Dis, 45, 985–1000.

31. Jang, H.J., Yang, Y.R., Kim, J.K., Choi, J.H., Seo, Y.K., Lee, Y.H., Lee, J.E., Ryu, S.H., & Suh, P.G. (2013). Phospholipase C-γ1 involved in brain disorders. Adv. Biol. Regul. 53, 51–62.

32. Kim, S.H., Yang, S., Lim, K.H., Ko, E., Jang, H.J., Kang, M., Suh, P.G., & Joo, J.Y. (2021). Prediction of Alzheimer’s disease-specific phospholipase c gamma-1 SNV by deep learning-based approach for high-throughput screening. Proc Natl Acad Sci U S A, 118, e2011250118.

33. Zou, C., Su, L., Pan, M., Chen, L., Li, H., Zou, C., Xie, J., Huang, X., Lu, M., & Zou, D. (2023). Exploration of novel biomarkers in Alzheimer’s disease based on four diagnostic models. Front Aging Neurosci, 15,1079433

34. Zhang, T., Liu, N., Wei, W., Zhang, Z., & Li, H. (2021). Integrated Analysis of Weighted Gene Coexpression Network Analysis Identifying Six Genes as Novel Biomarkers for Alzheimer’s Disease. Oxid Med Cell Longev, 2021, 9918498.

35. Wang, Y. & Liu, X. (2023). The Effective Components, Core Targets, and Key Pathways of Ginseng against Alzheimer’s Disease. Evid Based Complement Alternat Med, 2023, 9935942.

36. Huang, X.Y., Xue, L.L., Chen, T.B., Huangfu, L.R., Wang, T.H., Xiong, L.L., & Yu, C.Y. (2023). Miracle fruit seed as a potential supplement for the treatment of learning and memory disorders in Alzheimer’s disease. Front Pharmacol, 13, 1080753.

37. Yuan, Q., Wang, C.W., Shi, J., & Lin, Z.X. (2017). Effects of Ginkgo biloba on dementia: An overview of systematic reviews. J Ethnopharmacol, 195, 1–9.

38. Wang, J., Chen, X., Bai, W., Wang, Z., Xiao, W., & Zhu, J. (2021). Study on Mechanism of Ginkgo biloba L. Leaves for the Treatment of Neurodegenerative Diseases Based on Network Pharmacology. Neurochem Res, 46, 1881–1894.

39. Viña, J., Escudero, J., Baquero, M., Cebrián, M., Carbonell-Asíns, J.A., Muñoz, J.E., Satorres, E., Meléndez, J.C., Ferrer-Rebolleda, J., Cózar-Santiago, M.D.P., Santabárbara-Gómez, J.M., Jové, M., Pamplona, R., Tarazona-Santabalbina, F.J., & Borrás, C. (2022). Genistein effect on cognition in prodromal Alzheimer’s disease patients. The GENIAL clinical trial. Alzheimers Res Ther, 14, 164.

40. Zubair, T., & Bandyopadhyay, D. (2023). Small Molecule EGFR Inhibitors as Anti-Cancer Agents: Discovery, Mechanisms of Action, and Opportunities. Int J Mol Sci, 24, 2651.

41. Kurz, C., Walker, L., Rauchmann, B.S., & Perneczky, R. (2022). Dysfunction of the blood-brain barrier in Alzheimer’s disease: Evidence from human studies. Neuropathol Appl Neurobiol, 48, e12782.

42. Tournier, N., Goutal, S., Mairinger, S., Hernández-Lozano, I., Filip, T., Sauberer, M., Caillé, F., Breuil, L., Stanek, J., Freeman, A.F., Novarino, G., Truillet, C., Wanek, T., & Langer, O. (2021). Complete inhibition of ABCB1 and ABCG2 at the blood-brain barrier by co-infusion of erlotinib and tariquidar to improve brain delivery of the model ABCB1/ABCG2 substrate [11C]erlotinib. J Cereb Blood Flow Metab, 41, 1634–1646.

43. Ondaro, J., Hernandez-Eguiazu, H., Garciandia-Arcelus, M., Loera-Valencia, R., Rodriguez-Gómez, L., Jiménez-Zúñiga, A., Goikolea, J., Rodriguez-Rodriguez, P., Ruiz-Martinez, J., Moreno, F., Lopez de Munain, A., Holt, I.J., Gil-Bea, F.J., & Gereñu, G. (2022). Defects of Nutrient Signaling and Autophagy in Neurodegeneration. Front Cell Dev Biol, 10, 836196.

44. Johnson, C., Burkhart, D.L., & Haigis, K.M. (2022). Classification of KRAS-Activating Mutations and the Implications for Therapeutic Intervention. Cancer Discov, 12, 913–923.

45. Passaro, A., Jänne, P.A., Mok, T., & Peters, S. (2021). Overcoming therapy resistance in EGFR-mutant lung cancer. Nat Cancer, 2, 377–391.

