## Supplementary figures and images for "The MET growth signaling complex drives Alzheimer’s Disease-associated brain pathology in aged Shugoshin 1 mouse cohesinopathy model"

### Supplementary Fig 1A

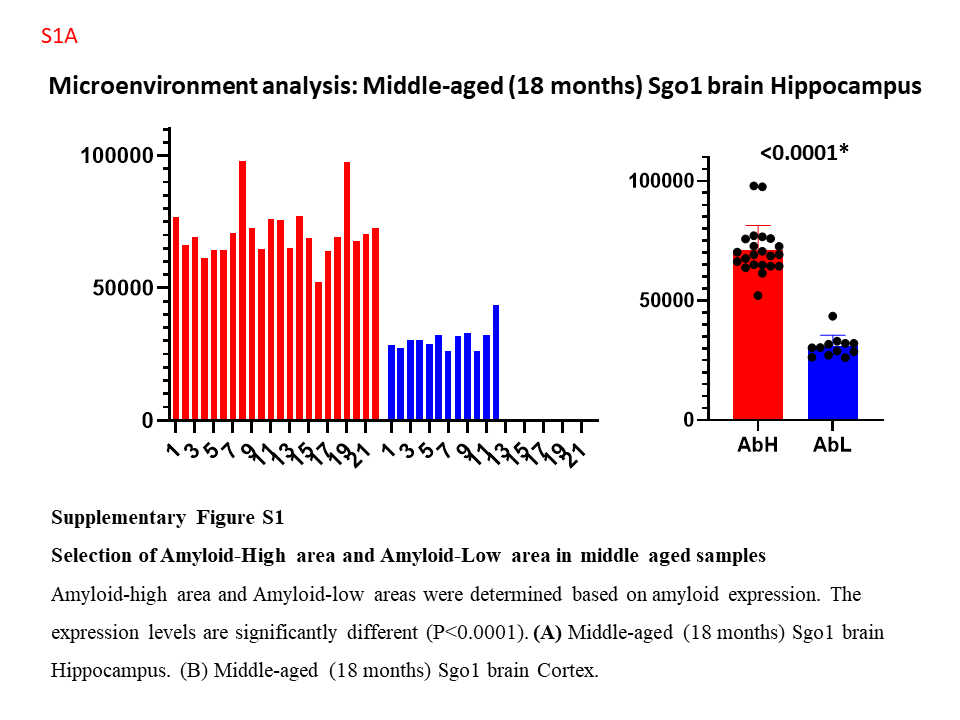

### Supplementary Fig 1B

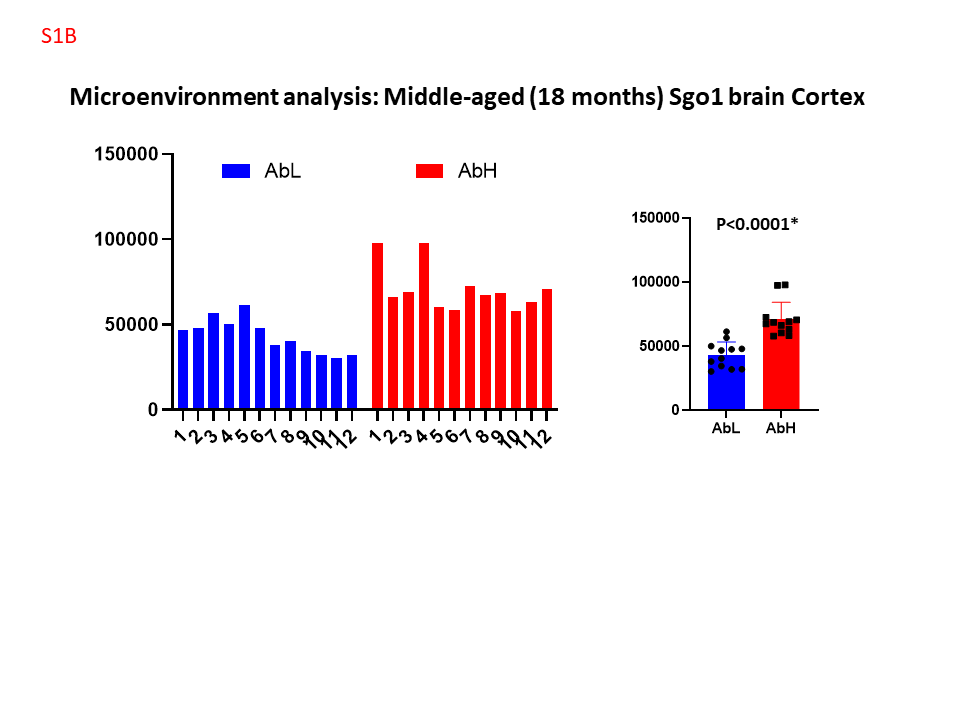

### Supplementary Fig 2

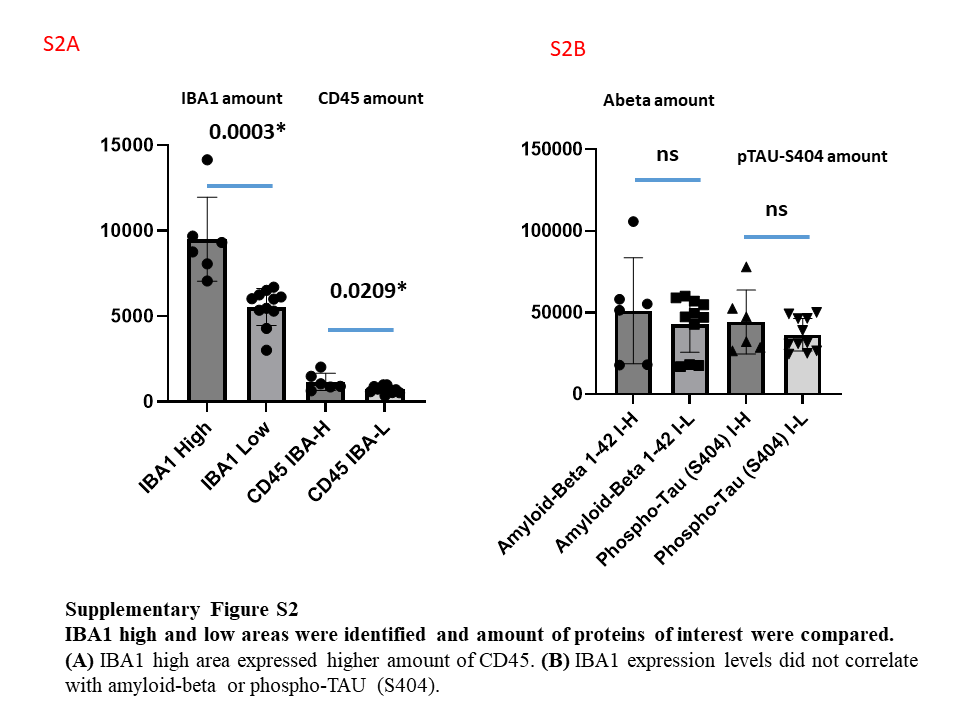
